# Multiple Omics Find New Cecal Microbial Features Associated with Feed Efficiency in Ducks

**DOI:** 10.1101/2024.04.25.591173

**Authors:** Rongbing Guo, Yuguang Chang, Dandan Wang, Hanxue Sun, Ayong Zhao, Tiantian Gu, Yibo Zong, Shiheng Zhou, Zhizhou Huang, Li Chen, Yong Tian, Wenwu Xu, Lizhi Lu, Tao Zeng

**Author notes:** Correspongding author at: State Key Laboratory for Managing Biotic and Chemical Threats to the Quality and Safety of Agro-products, Key Laboratory of Livestock and Poultry Resources (Poultry) Evaluation and Utilization, Ministry of Agriculture and Rural Affairs of China, Zhejiang Provincial Engineering Research Center for Poultry Breeding Industry and Green Farming Technology, Institute of Animal Science & Veterinary, Zhejiang Academy of Agricultural Sciences, Hangzhou 310021, China, Email Addresses (Lizhi Lu), (Tao Zeng).

## Abstract

As the global population continues to grow exponentially, the competition for resources between livestock and humans has become increasingly intense. Breeding efficient animal breeds, fully utilizing feed resources, and reducing environmental damage are major challenges facing the livestock industry. To address these issues, enhancing the feed utilization efficiency in the poultry industry is crucial. Recent studies have elucidated the pivotal role of gut microbiota in modulating the feeding behavior of their host organisms. Thus, we used metagenomics, transcriptomics, and metabolomics to explore how the intestinal microbiome affects the feed utilization efficiency in ducks. Our metagenomic analysis revealed a significant up-regulation of *Elusimicrobiota* at the phylum level within the high residual feed intake (HRFI) group, in comparison to the low residual feed intake (LRFI) group. Additionally, functional analysis using Clusters of Orthologous Groups of proteins (COG) indicated prominent disparities in the category of secondary metabolites biosynthesis, transport, and catabolism between the HRFI and LRFI groups. Furthermore, our metabolomics investigation identified an upregulated expression of the secondary metabolite 15-deoxy-Δ12,14-prostaglandin J2 (15d-PGJ2) in the HRFI group compared to the LRFI group. Liver transcriptome analysis identified prostaglandin-endoperoxide synthase 2 (*PTGS2*) as a key hub gene, exerting significant regulatory influence within the arachidonic acid pathway. Notably, the metabolite 15d-PGJ2 is a terminal product in the metabolic pathway of arachidonic acid. The correlation analysis between the cecal microbiota and differential metabolites revealed a significant negative correlation between *Elusimicrobiota* and the metabolite 15d-PGJ2. In summary, we assumed that the intestinal microbiome *Elusimicrobiota* regulates the expression of the *PTGS2* gene, consequently inducing variations in *PTGS2* efficiency between the HRFI and LRFI groups, ultimately leading to diverse residual feed intake levels in ducks.

**IMPORTANCE:** This investigation utilizes metabolomics to elucidate the interplay between genes and microbiome communities. We present evidence of disparities in the composition of microbial consortia among ducks RFI, alongside identification of pivotal genes within the liver that potentially modulate RFI. These results provide novel perspectives on the processes through which the cecum and liver influence RFI.

## INTRODUCTION

China is the world’s largest producer and consumer of duck. In addition to consuming duck meat and eggs, Chinese consumers also favor secondary products such as duck necks and wings (1). In recent years, the duck farming industry in China has undergone substantial and swift expansion. Nonetheless, such rapid advancements in livestock sectors have exerted considerable environmental strain. Additionally, the contention for resources between humans and livestock is escalating; therefore, our objective is to mitigate these issues through the propagation of efficient animal breeds. Residual feed intake (RFI), as proposed by Koch, serves as an index for estimating feed efficiency (2, 3). Animals with low RFI (LRFI) had higher feed use efficiency and animals with high RFI (HRFI) had lower feed use efficiency (4).

Animal feed conversion is closely related to lipid metabolism and energy metabolism (5). An increasing body of research has demonstrated that the gut microbiota exerts a significant influence on the feed efficiency of animals, particularly in the caecum (6–10). The liver is an indispensable and vital organ in humans and animals, serving as a central hub for various metabolic processes. These include protein metabolism, carbohydrate metabolism, lipid metabolism, vitamin metabolism, and hormone metabolism (11). The liver’s lipid metabolism products interact with the gut microbiota and other organs(12). Arachidonic acid (AA) and its derivative lipid mediators, such as prostaglandins (PGs), leukotrienes, and various other substances, play a crucial role in regulating hepatic lipid metabolism (13, 14). Recent research has uncovered a bidirectional communication axis between the gut and liver, which allows the gut microbiota to significantly affect an animal’s feeding behavior and energy metabolism (15). The various compounds produced within the intestinal tract can also influence the interactions between the gut microbiota and the host, such as short-chain fatty acids (16), bile acids (17), choline metabolites (18), amino acid-derived metabolites (19) and microbial components (20). These serve as signaling molecules that are detected by various host receptors, subsequently activating signaling and metabolic pathways in key tissues involved in energy metabolism and food intake regulation.

In recent years, the 16s rDNA sequencing technique has been one of the most commonly used methods for recognizing gut microbes. It mainly studies the species composition, and evolutionary relationship between species and community diversity (21–23). Metagenomics sequencing focuses on microbial population structure, gene function and activity, cooperation between microorganisms, and the relationship between microorganisms and the environment (24). Lots of experiments by LC-MS have suggested that it can accurately identify and quantify small molecules involved in metabolic reactions (25, 26). Besides it has excellent performance in the discovery of molecular markers(27). Some studies have combined 16S rDNA sequencing, metagenomes and metabolomics to better understand the composition, diversity, function and interaction mechanism of intestinal microorganisms (28). Over the past decade, transcriptome has become an increasingly mature technology. Nowadays, it can help us identify genes at the genetic level that underlie phenotypic differences between groups of samples (29).

Current research on the regulation of duck feeding behavior by the liver and microorganisms is limited. To deepen our understanding of the interaction between host genes and microbiota in ducks with varying residual feed intakes, we conducted an investigation integrating liver transcriptomics, caecum metagenomics, and metabolomics. Upon analyzing the composition and function of the intestinal microbial community, we have identified specific microbial species or metabolic products associated with the host’s genetic background. These findings are expected to serve as potential molecular markers for future genetic breeding initiatives.

## MATERIALS AND METHODS

### Animals and samples

Animal care and testing for this study were approved by the Institutional Animal Care and Use Committee of the Zhejiang Provincial Academy of Agricultural Sciences (License No.: 2022ZAASLA59) and were conducted in accordance with the regulations on the management of laboratory animals. In this experiment, 300 healthy 40-week-old ducks were selected. Their weights were recorded before and after the experiment, and the feed intake (FI) and egg mass loss (EML) of each duck were recorded daily. The RFI was calculated using the following formula:

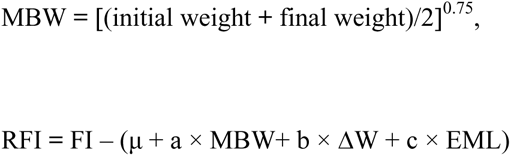

The experimental period was 5 weeks. The ducks were housed in a single cage to avoid any errors caused by pecking each other and were free to feed and drink, with the ambient temperature maintained at 15-27°C and the average light time at 16 h. The ducks were slaughtered at 45 weeks of age. The appendix contents and liver samples were stored at -80℃ until analysis.

### Metagenomic sequencing and data processing

Microbial DNA was extracted from the Shan Partridge duck samples using the E.Z.N.A.® Stool DNA Kit (Omega Bio-tek, Norcross, GA, U.S.) in accordance with the manufacturer’s protocols. Metagenomic shotgun sequencing libraries were constructed and sequenced at Shanghai Biozeron Biological Technology Co. Ltd. In summary, for each sample, 1μg of genomic DNA was sheared by a Covaris S220 Focused-ultrasonicator (Woburn, MA, USA), and sequencing libraries were prepared with a fragment length of approximately 450 bp. All samples were sequenced on an Illumina HiSeq instrument with paired-end 150bp (PE150) mode. Raw sequence reads underwent quality trimming using Trimmomatic to remove adaptor contaminants and low-quality reads (http://www.usadellab.org/cms/uploads/supplementary/Trimmomatic). Reads passing quality control were then mapped against the human genome (version: hg19) using the BWA mem algorithm (parameters: -M -k 32 -t 16; http://bio-bwa.sourceforge.net/bwa.shtml). The reads that removed host-genome contaminations and low-quality data were termed clean reads and used for further analysis.

### Untargeted Metabolomics Materials and Methods

The cecal contents (100 mg) were individually ground with liquid nitrogen, homogenized, and resuspended in cooled 80% methanol containing 0.1% formic acid. The samples were incubated on ice for 5 minutes and subsequently centrifuged at 15,000 rpm for 5 minutes at 4°C. A portion of the supernatant was diluted with LC-MS grade methanol to achieve a final concentration of 53% methanol. This mixture was then transferred to a new Eppendorf tube and centrifuged at 15,000 g for 10 minutes at 4°C. Finally, the supernatant was injected into the LC-MS system for analysis. UHPLC-MS/MS analyses were performed using a Vanquish UHPLC system (Thermo Fisher, Germany) coupled with an Orbitrap Q ExactiveTM HF mass spectrometer (Thermo Fisher, Germany) in Biozeron Co., Ltd. (Shanghai, China). The samples were injected onto a Hypesil Gold column (100×2.1 mm, 1.9μm) using a 17-minute linear gradient at a flow rate of 0.2 mL/min. The eluents for the positive polarity mode were eluent A (0.1% FA in Water) and eluent B (Methanol), while for the negative polarity mode, eluent A consisted of 5 mM ammonium acetate at pH 9.0 and eluent B was Methanol. The solvent gradient profile was set as follows: 2% B for 1.5 minutes; a gradient from 2% to 100% B over 12.0 minutes; 100% B for 2.0 minutes; a decrease from 100% to 2% B in 0.1 minutes; and finally, 2% B for the remaining 4.9 minutes.

### Transcriptomic sequencing and data processing

Total RNA was extracted from the liver tissue using TRIzol® Reagent according to the manufacturer’s instructions (Invitrogen) and genomic DNA was removed using DNase I (Takara). RNA-seq transcriptome libraries were prepared following the TruSeqTM RNA sample preparation Kit from Illumina (San Diego, CA), using 1μg of total RNA. After quantified by TBS380, paired-end libraries were sequenced by Illumina NovaSeq6000 sequencing (150bp*2, Shanghai BIOZERON Co., Ltd). The raw paired-end reads were trimmed and quality controlled by Trimmomatic with parameters (SLIDINGWINDOW:4:15 MINLEN:75) (version 0.36 http://www.usadellab.org/cms/uploads/supplementary/Trimmomatic). Then clean reads were separately aligned to Anas platyrhynchos reference genome with orientation mode using hisat2. Use htseq (https://htseq.readthedocs.io/en/release_0.11.1/) to count each gene reads. The expression levels of genes between the two groups were calculated using the fragments per kilobase of exon per million mapped reads (FRKM) method. R statistical package edgeR(Empirical Analysis of Digital Gene Expression in R, http://www.bioconductor.org/packages/release/bioc/html/edgeR.html/) was used to screen out differentially expressed genes (DEGs). When the logarithmic fold change was greater than 2 and the false discovery rate (FDR) should be less than 0.05 was considered as DEGs between the two groups. To understand the functions of the differentially expressed genes, GO functional enrichment and KEGG pathway analysis were carried out by Goatools (https://github.com/tanghaibao/Goatools) and KOBAS (http://kobas.cbi.pku.edu.cn/home.do). DEGs were significantly enriched in GO terms and metabolic pathways when their Bonferroni-corrected P-value was less than 0.05.

### Bioinformatics and statistical analysis

Co-occurrence among the bacterial taxa was analyzed using the SparCC program with the default settings. Spearman correlation analysis was performed to associate microbial taxa with the transcriptionally active functions (active functions hereafter). Only the genus-level bacterial taxa with a relative abundance > 0.1% and prevalence > 50% were used in the co-occurrence and correlation analysis, and only those with a correlation coefficient of > 0.5 or < −0.5 and a P value of < 0.05 were used in cooccurrence network analysis. Networks were visualized using Cytoscape (Version 3.9.1, http://www.cytoscape.org). The hubs of the microbes in the networks were calculated using the “CytoHubba” function in the Cytoscape software based on the Maximal Clique Centrality (MCC) method (https://apps.cytoscape.org/apps/cytohubba).

## RESULTS AND DISCUSSION

### Animal phenotypes data analysis

To compare the HRFI group and LRFI group body weight (BW) and the daily egg mass (EML) were similar between the HRFI and the LRFI ducks (*P* > 0.05), but daily feed intake (FI), residual feed intake (RFI) and feed conversion ratio (FCR) were higher (*P* < 0.01) in the HRFI ducks than in the LRFI ducks (Fig.1A).

**Fig. 1.**
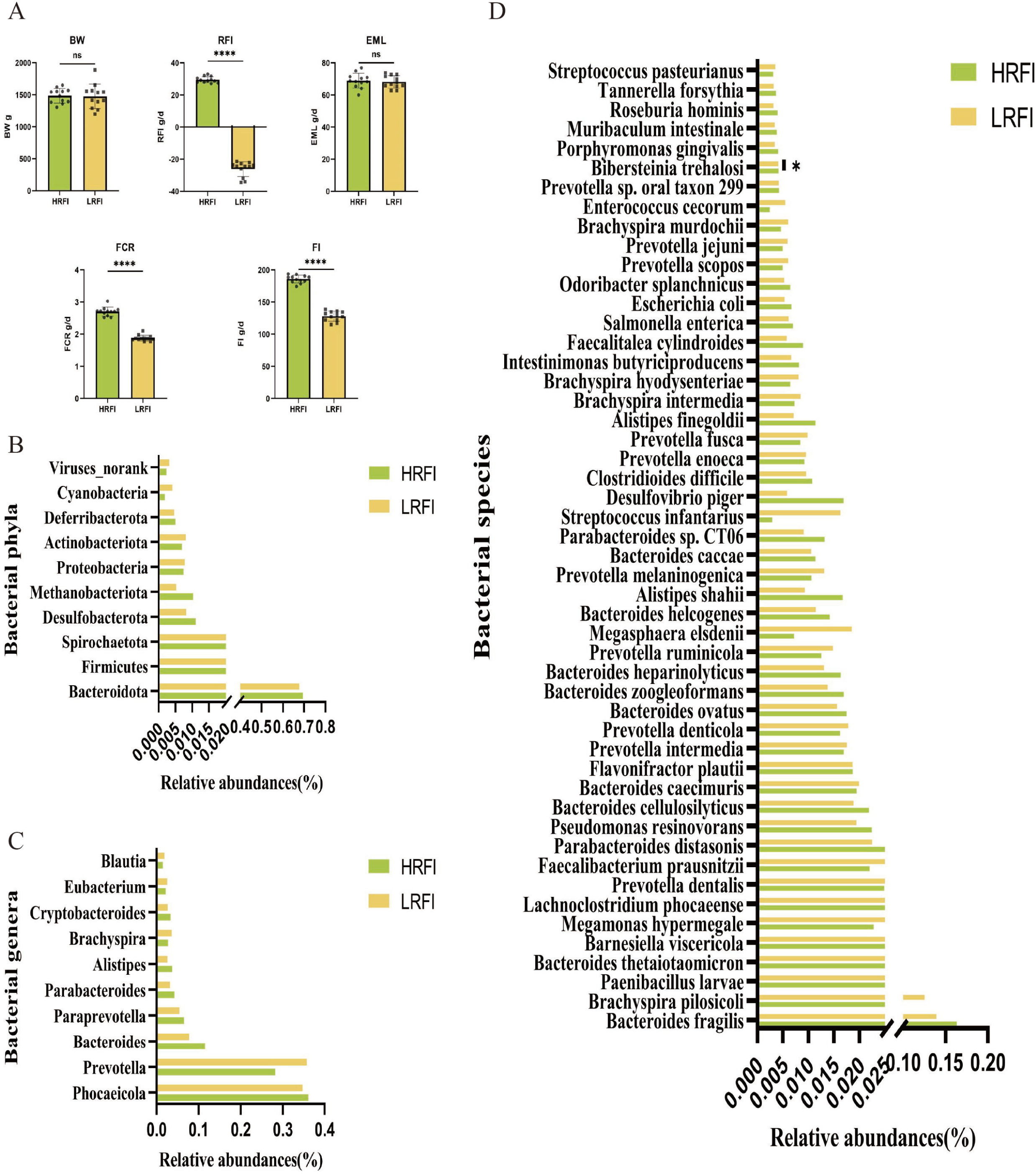
Comparison of phenotypic data and cecum bacterial taxa identified in the metagenomes between ducks with different feed efficiency. Body weight (BW), Residual feed intake(RFI), the daily egg mass(EML), Feed conversion rate (FCR) and actual feed intake(FI) were compared using a t test (A). The 10 most abundant bacterial phyla (B), 10 most abundant bacterial genera (C), and 50 most abundant bacterial species (D). The T-test was used for mean comparison. *P < 0.05

### Metagenomic sequencing data analysis

A total of 163 Gb of data were obtained from the metagenomic sequencing, with 10.21 ± 1.82 GB per sample (Supplementary TableS1). A total of 137.16 GB of data was retained after quality filtering and removing host DNA sequences. A total of 3,889,047 contigs were generated from de novo assembly (486,131 ± 83,703 per sample, N50 length of 2,109 ± 210).

Prom the metagenomic sequences of bacteria (11,772,155 ± 2,499,903 sequences per sample), a total of 34 phyla, 1033 genera, and 3262 species of bacteria were identified (data not shown). At the phylum level, we select the mean relative abundance > 0.1% microbiomes. We found that *Elusimicrobiota* was more abundant (*P* < 0.05) in the HRFI ducks than in the LRFI ducks (Fig.1B). As for the top10 abundance of genus level, we found that none of these predominant bacterial genera differed in relative abundance between the two duck groups (Fig.1C). At the species level, we selected the top50 abundance microbiomes, and found a species called *Bibersteinia trehalosi (B. trehalosi)* had a higher abundance (*P* < 0.05) in the LRFI ducks than in the HRFI ducks (Fig.1D, Supplement TableS2).

The metagenomic sequencing mapped a total of 5 Kyoto Encyclopedia of Genes and Genomes (KEGG) level-1 pathways, 34 level-2 KEGG pathways, and 321 KEGG level-3 pathways. In the level-3 KEGG pathway, the “Metabolism” (47.66%), “Genetic Information Processing” (6.85%), “Environmental Information Processing” (10.90%), “Cellular Processes” (9.66%) and “Organismal Systems” (24.92%). The level-3 KEGG pathway of abundance top20 was compared and no difference was found between the two groups of HRFI ducks and LRFI ducks (Supplementary TableS3, Fig.2A).

**Fig. 2.**
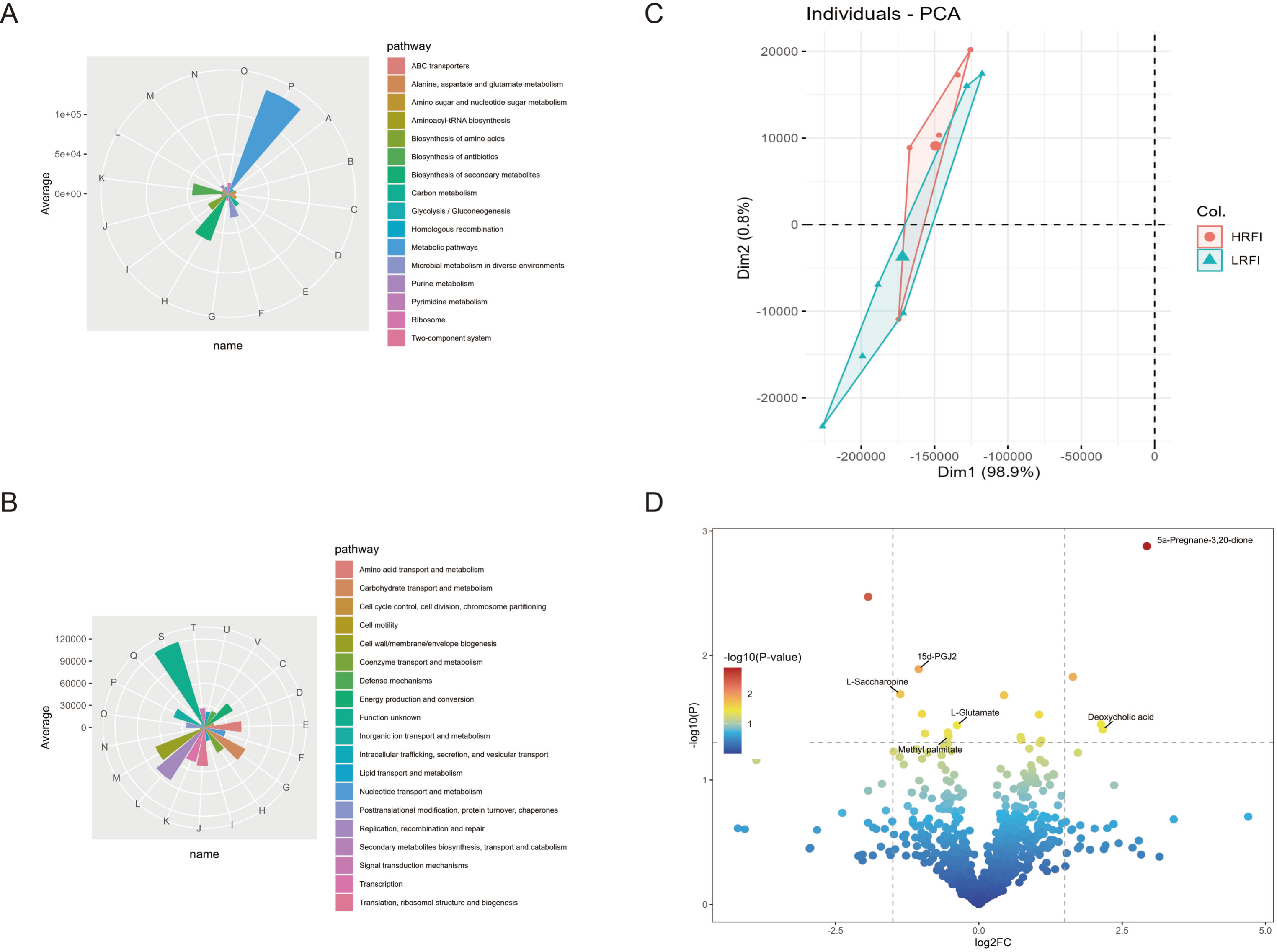
Metagenomic KEGG and COG pathway map and metabolomics data analysis. The different colors in the figure represent different pathways, and the size of the color blocks in the figure represents the average abundance (A). Metagenomic COG pathway map. The different colors in the figure represent different pathways, and the size of the color blocks in the figure represents the average abundance (B). PCA analysis of HRFI and LRFI duck metabolites (C). HRFI and LRFI ducks metabolites expression. A curated list of 922 metabolites of the cecum were analyzed the associated with possible effects on feed intake genes were labeled on the plot. Two vertical lines indicate gene expression fold change (HRFI vs. LRFI) >1.2 and <−1.2, respectively, and the horizontal line indicates the adjusted P value (FDR q-value) of 0.05. P values were calculated by two-sided Wilcoxon rank-sum test. The color of the dot represents the FDR (q-value) levels (D).

Clusters of Orthologous Groups of proteins (COG) have been a popular tool in microbial genome annotation. In our study, a total of 24 functional categories were enriched. Among them, the Secondary metabolites biosynthesis, transport and catabolism category showed significant differences between HRFI and LRFI groups (*P* < 0.05, Fig.2B).

### The gut microbiota mainly affects Lipids and lipid like molecules metabolism

To identify key metabolites regulated by intestinal commensal bacteria that may affect duck feeding, we performed LC-MS non-targeted metabolomics analysis of the cecum contents of the mountain mallard duck, and a total of 922 metabolites were identified. The compositions between the two groups were very similar to the PCA plots of the differences between the two groups (Fig 2C), in which there were 17 metabolites with significant differences between the two groups. From the differential metabolites, we found that 15-Deoxy-Delta-12,14-Prostaglandin J2 (15d-PGJ2), L Saccharopine, L Glutamate, Mehty Palmitate, and Deoxycholic acid are associated with the physiological processes of fat digestion, absorption, and synthesis (Fig.2D). KEGG pathway of differential metabolites were primarily enriched in Lipids and lipid-like molecules, Organic acids and derivatives, Benzenoids, Organoheterocyclic compounds and Phenylpropanoids and polketides Super Class (HMDB). Among the differential metabolites, the most abundant one was 2-Hydroxycinnamic acid.

### The gut microbiota regulates the expression of liver genes involved in duck fat digestion and absorption

The average total numbers of raw reads and raw bases in the sample were approximately 100 million and 16 billion, respectively. The average total numbers of clean reads and clean bases in the sample were approximately 100 million and 15 billion, respectively. The percent of raw reads and raw bases respectively were 99.3% and 96.7%. The average GC content of the samples was approximately 49%, whereas the average percentages of Q20 and Q30 bases were 99.4% and 97.6% (Supplement TableS5).

The PCA can reflect the overall expression differences between groups and the degree of variation within samples. Our analysis results show that the samples clustered between groups, indicating no significant difference between groups (Fig.3A). Then we explored changes in the duck liver transcriptome induced by intestinal microbiota. RNA sequencing analysis revealed that 322 genes had differential expression (log_2_FC > 2, FDR < 0.05, Fig.3B). The liver is the metabolic center within an animal’s body, and it is closely associated with the animal’s feeding behavior (30). We found that genes encoding fatty acid-binding protein (*FABP1*, *FABP3*) in the liver can convert arachidonic acid to prostaglandin endoperoxide synthase *PTGS2*, and genes related to lipid digestion and transport, secreted by the pancreas, are crucial for lipid metabolism. By comparing the HRFI group and the LRFI group, we found that *FABP1* and *FABP3* showed a downward trend, while *PTGS*2 showed an upward trend. KEGG enrichment analysis of differentially expressed genes revealed that Fat digestion and absorption in the Digestive system were significantly different between the two groups (*P* < 0.05, Fig 3E).

**Fig. 3.**
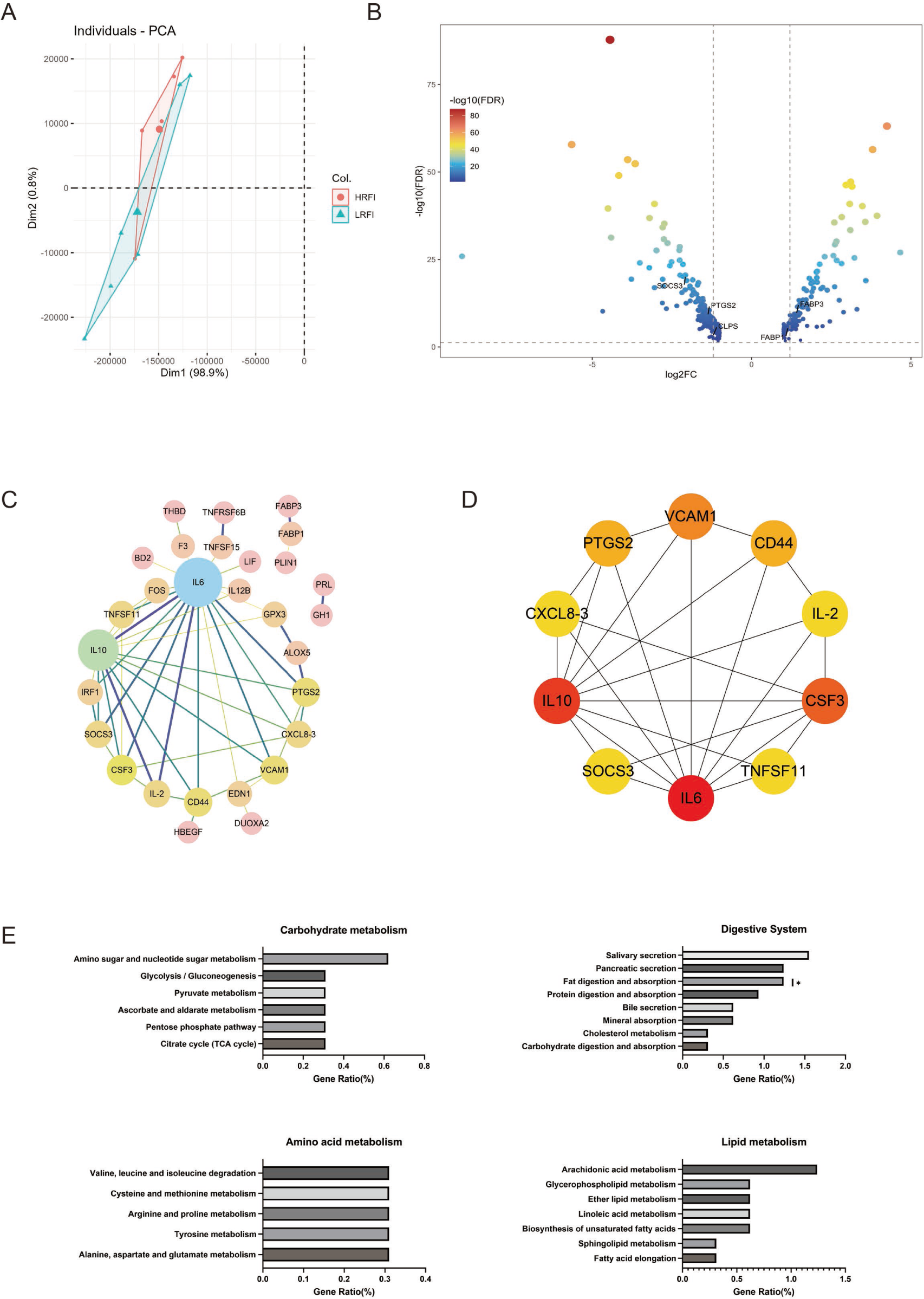
Transcriptome data analysis. PCA analysis of HRFI and LRFI duck genes (A). Differentially expressed liver genes between HRFI and LRFI ducks. A curated list of 338 genes of the liver were analyzed the associated with possible effects on feed intake genes were labeled on the plot. Two vertical lines indicate gene expression fold change (HRFI vs. LRFI) >1.2 and <−1.2, respectively, and the horizontal line indicates the adjusted P value (FDR q-value) of 0.05. P values were calculated by two-sided Wilcoxon rank-sum test. The color of the dot represents the FDR (q-value) levels (B). PPI network diagram of liver differential genes. Nodes represent proteins. The node size and color represent the mean abundance of genes expression. Edges represent protein-protein associations. The relationship between the 2 proteins is expressed through the thickness of the line; the thicker the line, the closer the relationship. The color represents the combined score, it was analysed by cytoscape (C). Hub genes and expression profiles of PPI net work. Degree is used as the evaluation criterion, and the darker the color of the node, the higher its Degree score (D). Pathways identified in the transcriptomes. The Wilcoxon rank-sum test was used for mean comparison, *P < 0.05 (E).

### Analysis of PPI Network for DEGs

To identify key genes associated with animal feeding behavior, we performed a PPI network analysis of the differentially expressed genes in significantly different pathways between the two groups. We finally obtained 29 nodes and 50 edges (Fig.3C). Degree represents the number of connections to that node. From this network diagram, we can observe that Interleukin 6 (*IL6*) has the most connections (Degree = 18), while the second bromodomain (*BD2*) has the least (Degree = 1,). To identify the core genes, we performed Cytohubba analysis on this PPI network and applied the MCC score method to select the top 10 most important core genes. The interleukin genes Interleukin 6 (*IL6*, Degree = 70) and interleukin 10 (*IL10*, Degree = 64) are located at the first two positions of the core genes. The *PTGS2* gene is listed at the fifth position in the core genes (Degree = 13, Fig.3D, Supplement tableS6).

### Key Metabolites, Microbiome and Metabolic Pathways co-occurrence network

To further understand which specific bacteria are involved in regulating RFI, we conducted a correlation analysis between the microbial communities in the metagenomics and the differential metabolites in the metabolic profile. We conducted a correlation analysis between the top 30 abundant microbial species in the metagenomics, KEGG pathways with significant differences, and five important metabolites.

Between metabolites and the top 10 KEGG pathways, there are a total of 50 interactive edges. Among them, Amino sugar and nucleotide sugar metabolism has a negative correlation with Deoxycholic acid (Pearson Correlation Coefficient (PCC) = -0.65, p = 0.08), while the metabolite L Glutamate and Methyl palmitate has a positive correlation with the Amino sugar and nucleotide sugar metabolism pathway, with PCC values of 0.61 and 0.60, respectively, and P values were 0.11 and 0.33. As for the key differential metabolites interacting with the microbial genus level, there are a total of 145 interactive edges. The highest significant positive correlation is that between *Wallbacteria* and Methyl palmitate (PCC = 0.93, *P* < 0.01), while the most significant negatively correlated relationship is that between *Parabacteroides* and L Glutamate (PCC =-0.92, *P* < 0.01, Fig.4A).

**Fig. 4.**
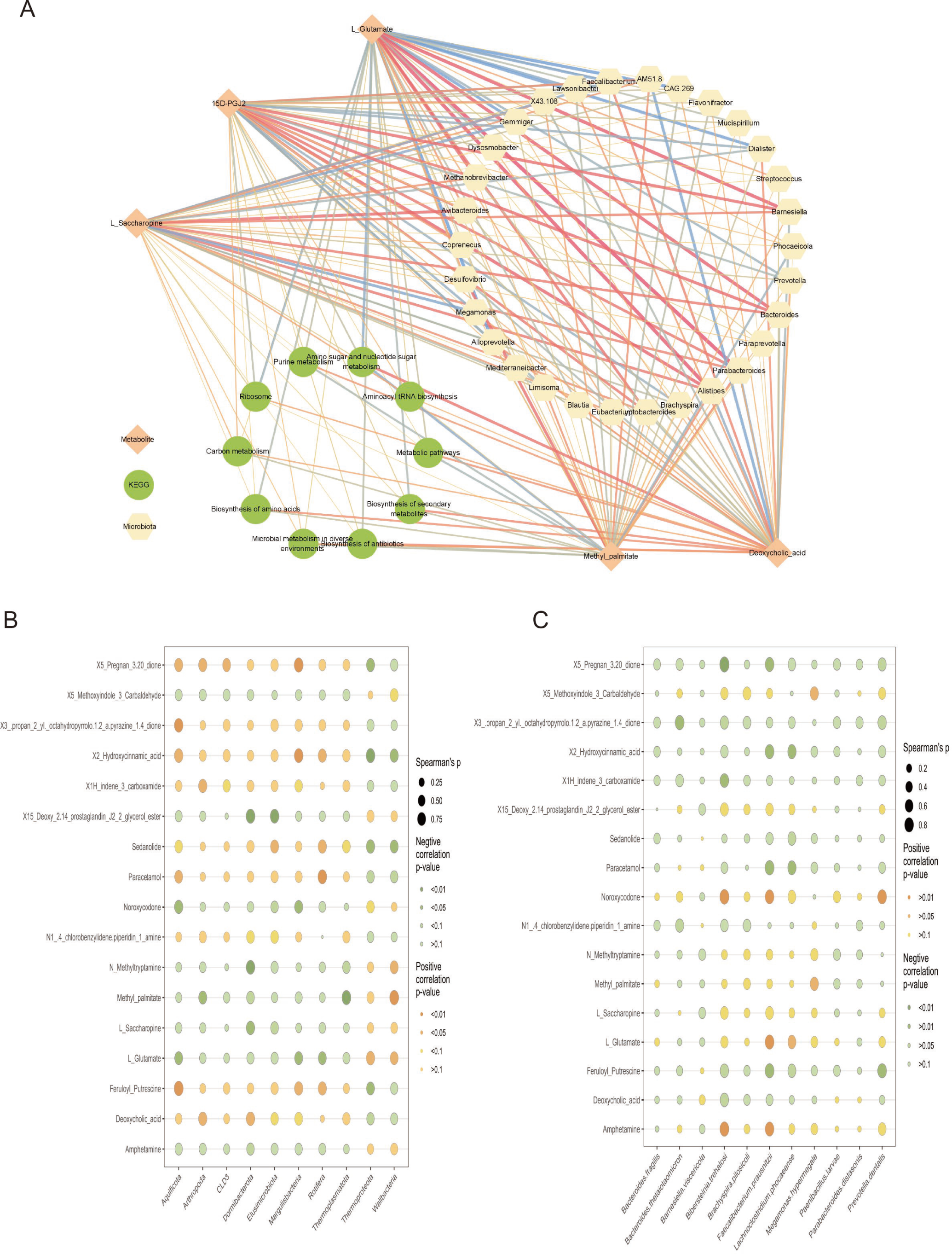
Co-occurrence network and data association analysis. The co-occurrence network among caecum bacteria and metabolites in ducks with high and low feed residual intake. Relationships between caecum genus level top30 abundance microbial, significantly different top10 microbial functions and 5 differential metabolites related to fat metabolism. Blue edges indicate positive relationships, and red edges indicate negative relationships (A). Correlation Analysis between Microbial Community Levels and Metabolic Products in the Cecum. We ranked the microorganisms at the phylum level from small to large, selected the top 10 phylum level bacteria with 17 differential metabolic products, and performed correlation analysis between the top 10 abundance of species level and the top 50. *B. trehalosi*, which showed significant differences between the two groups. The bubble chart was plotted using the calculated log2 fold change (HRFI vs. LRFI) and p values. The size of the bubble indicates the statistical difference, with larger bubbles indicating more significant correlations. The color of the bubble represents the positive or negative correlation between the microorganisms and differential metabolic products. Orange represents positive correlation, while green represents negative correlation (B).

Firstly, at the level of bacterial phyla, the *Eulisimicrobiota* showed a highly significant negative correlation with the metabolite 15-Deoxy-Delta-12,14-Prostaglandin J2 (15d-PGJ2, cor = -0.86, *P* < 0.01, Fig.4B). There was a significant positive correlation between *Sedanolide* and the metabolite (cor = 0.74, *P* < 0.05). For the metabolite Deoxycholic acid, there was a significant positive correlation between *Dormibacterota* and Arthropoda (*P* < 0.05), with a correlation coefficient of 0.80 and 0.78 respectively. There was also a significant negative correlation between 15d-PGJ2 and *Dormibacterota* (cor = -0.89, *P* < 0.05). the level of bacterial species, our study results showed that *B. trehalosi* had a significant positive correlation with metabolites Noroxycodone and amphetamine, with correlation coefficients of 0.80 and 0.76 respectively (*P* < 0.05, Fig.4C). *B. trehalosi* had a highly significant negative correlation with the metabolite X5 Pregnan-3.20-dione (cor = -0.86, *P* < 0.01).

## Discussion

Feed efficiency is a critical factor in reducing the costs associated with livestock production and enhancing environmental protection (31). RFI serves as a commonly employed metric for assessing feed efficiency. It is calculated by regressing the feed intake against the EML, BW and the BW^0.75^ (32, 33). Research indicates that RFI may be connected to mechanisms related to feeding and digestion (34). In recent years, an increasing number of scholars have recognized the pivotal role of the gut microbiota in the digestion and absorption of nutrients from animal feed, which is also likely to influence the efficiency of feed utilization in animals (35–38).

Currently, the two predominant techniques utilized for acquiring knowledge about gut microorganisms are 16S rRNA sequencing and metagenomic analysis. While 16S rRNA sequencing primarily elucidates the species composition within communities, delineates evolutionary relationships among them, and assesses their diversity, metagenomic sequencing facilitates comprehensive investigations into genetic and functional aspects based on the preliminary insights gained from 16S rRNA data (39, 40). In our investigation, we employed a comprehensive metagenomic approach to analyze the microbial composition and functional attributes of the microorganisms present in the cecal contents of ducks.

Our findings indicated that, although the HRFI and LRFI groups exhibited considerable similarity in their overall microbial colony composition, significant differences were observed at the level of specific colonies. In terms of phylum levels, the top two most abundant phyla were the *Bacteroidota* and *Firmicutes* phylum, respectively, and the expression of *Elusimicrobiota* was significantly different between two groups. Liu et al.’s study found that the expression of *Elusimicrobiota* in yaks drinking warm water with better growth performance was significantly lower than that in yaks drinking cold water with poorer growth performance (41). Our findings indicate that the *Elusimicrobiota* population is notably higher in the HRFI group compared to the LRFI group of ducks, with a statistically significant difference (*P* = 0.04). This phenomenon suggests that the microbe *Elusimicrobiota* may also influence the productive performance of ducks.

The liver serves as the metabolic hub in animals, undertaking a multitude of functions including lipid metabolism, protein metabolism, carbohydrate metabolism, bile secretion, detoxification, and immune defense (42, 43). Fatty acids are not merely precursors to numerous vital bioactive molecules, such as prostacyclin, prostaglandins, and leukotrienes (44–46), but also constitute an essential energy source integral to diverse biosynthetic processes within organisms (47). Studies have shown that the process of fatty acid oxidation can elicit feeding behavior in rats. (48). *PTGS2*, also recognized as Cyclooxygenase-2 (*COX-2*), is not only a crucial enzyme implicated in arachidonic acid metabolism (49) but also the pivotal catalyst for the rate-limiting step in the transformation of arachidonic acid into prostaglandins (50). The inherent expression of human *COX-2* (*hCOX-2*) within hepatocytes may forestall obesity induced by high-fat diets by stimulating thermogenesis and fatty acid oxidation (51). *COX-2* facilitates the synthesis of arachidonic acid to yield PGD2 (52). As a dehydrated variant of PGD2, it can also modulate inflammatory response and immune system functions by suppressing the generation of other prostaglandins (53). In our study of the liver transcriptome, among the KEGG pathways enriched within the digestive system, the fat digestion and absorption pathway exhibited notable differences between groups. Further, the expression of *PTGS2* was significantly lower in the LRFI group compared to the HRFI group. Interestingly, the differential metabolite 15d-PGJ2 was significantly higher in the LRFI group compared to the HRFI group. Based on our observations, we hypothesize that the LRFI group exhibits higher utilization efficiency of *PTGS2* compared to the HRFI group, leading to an increased expression of 15d-PGJ2 in the LRFI group. It is noteworthy that upon analyzing the metabolic pathways where the differential metabolites were located, we discovered that the key gene *PTGS2* and the significantly different metabolite 15d-PGJ2 were both enriched in the arachidonic acid pathway (Fig. 5A). Furthermore, research has indicated a potential association between the metabolism of arachidonic acid and the RFI phenotype. (54). Although there was no significant difference in the enrichment of Arachidonic acid metabolism pathways between the two groups in our transcriptome results (*P* = 0.08), based on the analysis of genes and metabolites mentioned above, we cannot deny its important role in regulating the phenotype of RFI. In summary, our study speculates that the microbiome *Elusimicrobiota*, may influence RFI variability in ducks by regulating the expression of the liver gene *PTGS2* (Fig. 5B).

**Fig. 5.**
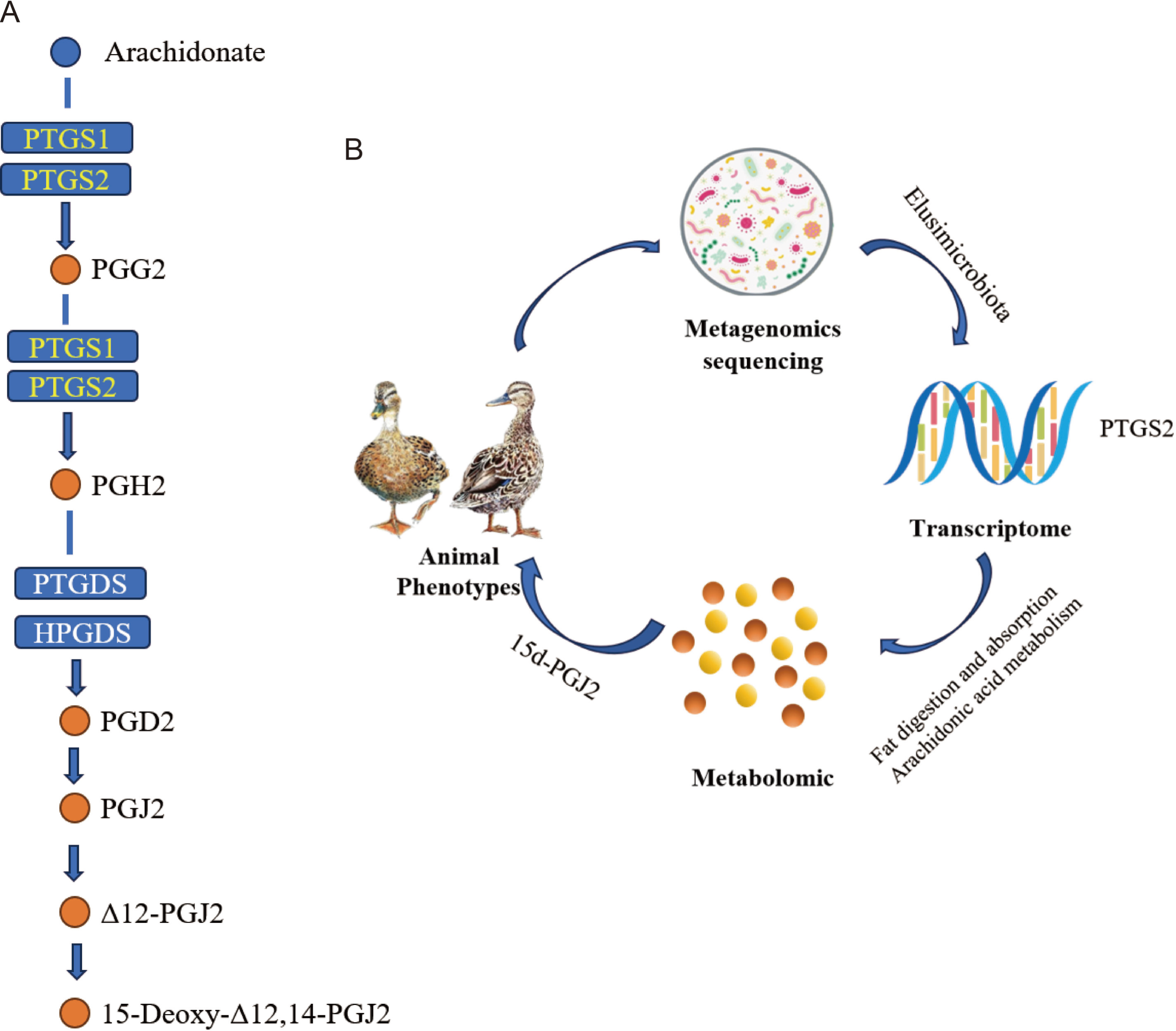
Arachidonic acid pathway and pathway regulation mechanism diagram. The hub gene *PTGS2* and differential metabolite 15d-PGJ2 were enriched in arachidonic acid metabolism pathway (A). Graphical summary of effects of intestinal microbiota on feeding behavior and phenotype in ducks (B).

## Conclusions

Our research findings offer valuable insights into interventions at the genetic, microbial, and metabolite levels that can enhance animal feed efficiency. These discoveries are crucial for improving animal feed efficiency and reducing competition for resources between humans and livestock. Our study suggests a complex interplay between the gut microbiome and the liver transcriptome. However, further experiments are required to validate our results and elucidate the specific underlying molecular mechanisms.

## ACKNOWLEDGMENTS

This study was supported by Zhejiang Provincial Natural Science Foundation of China (Grant No. LZ23C170001), the National Key Research and Development Program of China (2022YFD1300100), China Agriculture Research System of MOF and MARA (CARS-42), and Zhejiang Province Agricultural New Breeding Major Science and Technology Special Project (2021C02068).

We also acknowledge the staff of Cherry Valley Agricultural Technology Co., Ltd. for their technical work on the experiments.

## Author Statement

All authors undersigned declare that this manuscript is original, has not been published before and is not currently being considered for publication elsewhere. We confirm that the manuscript has been read and approved by all named authors and that there are no other persons who satisfied the criteria for authorship but are not listed.

## CRediT authorship contribution statement

**Rongbing Guo:** Formal analysis, Investigation, Methodology, Visualization, Writing – original draft, Writing – review & editing. **Yuguang Chang:** Data curation, Formal analysis, Methodology, Software. **Dandan Wang:** Investigation. **Hanxue Sun** and **Ayong Zhao:** Samples collection. **Tiantian Gu and Yibo Zong:** Writing – review & editing. **Shiheng Zhou and Zhizhou Huang:** Provide us with the animal experiment site. **Li Chen, Yong Tian and Wenwu Xu:** Offering writing suggestions. **Lizhi Lu:** Resources. **Tao Zeng:** Conceptualization, Formal analysis, Funding acquisition, Methodology, Supervision, Writing – review & editing.

## Data availability

The metagenomic and transcriptomic raw data have been deposited in the CNGB Sequence Archive (CNSA) of China National Gene Bank Data Base (CNGBdb) under the accession numbers CRA013513 and CRA014753, respectively.

## References

1. Zhang F, Zhu F, Yang FX, Hao JP, Hou ZC. 2022. Genomic selection for meat quality traits in Pekin duck. Anim Genet 53:94–100.10.1111/age.13157.

2. Barendse W, Reverter A, Bunch RJ, Harrison BE, Barris W, Thomas MB. 2007. A validated whole-genome association study of efficient food conversion in cattle. Genetics 176:1893–905.10.1534/genetics.107.072637.

3. Koch RM GK, Chambers D, Swiger LA. 1963. Efficiency of feed use in beef cattle. . J Anim Sci.

4. Patience JF, Rossoni-Serao MC, Gutierrez NA. 2015. A review of feed efficiency in swine: biology and application. J Anim Sci Biotechnol 6:33.10.1186/s40104-015-0031-2.

5. Patience JF, Rossoni-Serão MC, NA G. 2015. A review of feed efficiency in swine: biology and application. . J Anim Sci Biotechnol doi:10.1186/s40104-015-0031-2.:6(1):33.10.1186/s40104-015-0031-2.

6. Xue MY, Xie YY, Zhong Y, Ma XJ, Sun HZ, Liu JX. 2022. Integrated meta-omics reveals new ruminal microbial features associated with feed efficiency in dairy cattle. Microbiome 10:32.10.1186/s40168-022-01228-9.

7. Yang H, Huang X, Fang S, He M, Zhao Y, Wu Z, Yang M, Zhang Z, Chen C, Huang L. 2017. Unraveling the Fecal Microbiota and Metagenomic Functional Capacity Associated with Feed Efficiency in Pigs. Front Microbiol 8:1555.10.3389/fmicb.2017.01555.

8. Bai H, Shi L, Guo Q, Jiang Y, Li X, Geng D, Wang C, Bi Y, Wang Z, Chen G, Xue F, Chang G. 2022. Metagenomic insights into the relationship between gut microbiota and residual feed intake of small-sized meat ducks. Front Microbiol 13:1075610.10.3389/fmicb.2022.1075610.

9. Wen C, Yan W, Mai C, Duan Z, Zheng J, Sun C, Yang N. 2021. Joint contributions of the gut microbiota and host genetics to feed efficiency in chickens. Microbiome 9:126.10.1186/s40168-021-01040-x.

10. Liu Y, Wu H, Chen W, Liu C, Meng Q, Zhou Z. 2022. Rumen Microbiome and Metabolome of High and Low Residual Feed Intake Angus Heifers. Front Vet Sci 9:812861.10.3389/fvets.2022.812861.

11. Trefts E, Gannon M, Wasserman DH. 2017. The liver. Current Biology 27:R1147–R1151.10.1016/j.cub.2017.09.019.

12. Canbay A, Bechmann L, Gerken G. 2007. Lipid Metabolism in the Liver. Zeitschrift für Gastroenterologie 45:35–41.10.1055/s-2006-927368.

13. Sarah A. Martin, Alan R. Brash, Murphy RC. 2016. The discovery and early structural studies of arachidonic acid. Journal of Lipid Research 57:1126–1132.10.1194/jlr.R068072.

14. Zhou Y, Khan H, Xiao J, WS. C. 2021. Effects of Arachidonic Acid Metabolites on Cardiovascular Health and Disease. . Int J Mol Sci doi:10.3390/ijms222112029.:22(21):12029.10.3390/ijms222112029.

15. Ringseis R, Gessner DK, Eder K. 2020. The Gut–Liver Axis in the Control of Energy Metabolism and Food Intake in Animals. Annual Review of Animal Biosciences 8:295–319.10.1146/annurev-animal-021419-083852.

16. Kindt A, Liebisch G, Clavel T, Haller D, Hörmannsperger G, Yoon H, Kolmeder D, Sigruener A, Krautbauer S, Seeliger C, Ganzha A, Schweizer S, Morisset R, Strowig T, Daniel H, Helm D, Küster B, Krumsiek J, Ecker J. 2018. The gut microbiota promotes hepatic fatty acid desaturation and elongation in mice. Nature Communications 9.10.1038/s41467-018-05767-4.

17. Trauner M, JL B. 2003. Bile salt transporters: molecular characterization, function, and regulation. Physiol Rev 2:633–71.10.1152/physrev.00027.2002.

18. Koeth RA, Wang Z, Levison BS, Buffa JA, Org E, Sheehy BT, Britt EB, Fu X, Wu Y, Li L, Smith JD, DiDonato JA, Chen J, Li H, Wu GD, Lewis JD, Warrier M, Brown JM, Krauss RM, Tang WHW, Bushman FD, Lusis AJ, Hazen SL. 2013. Intestinal microbiota metabolism of l-carnitine, a nutrient in red meat, promotes atherosclerosis. Nature Medicine 19:576–585.10.1038/nm.3145.

19. Beaumont M, Neyrinck AM, Olivares M, Rodriguez J, de Rocca Serra A, Roumain M, Bindels LB, Cani PD, Evenepoel P, Muccioli GG, Demoulin JB, NM. D. 2018. The gut microbiota metabolite indole alleviates liver inflammation in mice. FASEB J 12:fj201800544.10.1096/fj.201800544.

20. Jenne CN, P K. 2013. Immune surveillance by the liver. Nat Immunol 10:996–1006.10.1038/ni.2691.

21. Shi DW, Wang DM, Ning LH, Li J, Dong Y, Zhang ZK, Dou HW, Wan RJ, Jia CM, L. X. 2022. Using 16S rDNA Sequencing Technology to Preliminarily Analyze Intestinal Flora in Children with Mycoplasma pneumoniae Pneumonia. . Biomed Environ Sci doi:10.3967/bes2022.070.:35(6):528-537.10.3967/bes2022.070.

22. Laudadio I, Fulci V, Palone F, Stronati L, Cucchiara S, C C. 2018. Quantitative Assessment of Shotgun Metagenomics and 16S rDNA Amplicon Sequencing in the Study of Human Gut Microbiome. . OMICS doi:10.1089/omi.2018.0013:22(4):248-254.10.1089/omi.2018.0013.

23. Kim H, Kim S, S. J. 2020. Instruction of microbiome taxonomic profiling based on 16S rRNA sequencing. . J Microbiol doi:10.1007/s12275-020-9556-y.:58(3):193-205.10.1007/s12275-020-9556-y.

24. Shi Y, Wang G, Lau HC, J. Y. 2022. Metagenomic Sequencing for Microbial DNA in Human Samples: Emerging Technological Advances. Int J Mol Sci doi:10.3390/ijms23042181. :23(4):2181.10.3390/ijms23042181. .

25. Zhou B, Xiao JF, Tuli L, Ressom HW. 2012. LC-MS-based metabolomics. Mol Biosyst 8:470–81.10.1039/c1mb05350g.

26. Artati A, Prehn C, Adamski J. 2019. LC-MS/MS-Based Metabolomics for Cell Cultures, p 119-130, Cell-Based Assays Using iPSCs for Drug Development and Testing doi:10.1007/978-1-4939-9477-9_10.

27. Chen CJ, Lee DY, Yu J, Lin YN, Lin TM. 2022. Recent advances in LC-MS-based metabolomics for clinical biomarker discovery. Mass Spectrom Rev doi:10.1002/mas.21785:e21785.10.1002/mas.21785.

28. Thingholm LB, Ruhlemann MC, Koch M, Fuqua B, Laucke G, Boehm R, Bang C, Franzosa EA, Hubenthal M, Rahnavard A, Frost F, Lloyd-Price J, Schirmer M, Lusis AJ, Vulpe CD, Lerch MM, Homuth G, Kacprowski T, Schmidt CO, Nothlings U, Karlsen TH, Lieb W, Laudes M, Franke A, Huttenhower C. 2019. Obese Individuals with and without Type 2 Diabetes Show Different Gut Microbial Functional Capacity and Composition. Cell Host Microbe 26:252–264 e10.10.1016/j.chom.2019.07.004.

29. Garcia-Nieto PE, Wang B, Fraser HB. 2022. Transcriptome diversity is a systematic source of variation in RNA-sequencing data. PLoS Comput Biol 18:e1009939.10.1371/journal.pcbi.1009939.

30. JM F. 1982. The role of the liver in the control of food intake. . Proc Nutr Soc doi:10.1079/pns19820021:41(2):123-6. .10.1079/pns19820021.

31. Touitou F, Tortereau F, Bret L, Marty-Gasset N, Marcon D, A M. 2022. Evaluation of the Links between Lamb Feed Efficiency and Rumen and Plasma Metabolomic Data. Metabolites doi:10.3390/metabo12040304:12(4):304.10.3390/metabo12040304 .

32. P Luiting, Urff. EM. 1991. Optimization of a model to estimate residual feed consumption in the laying hen. Livestock Production Science 10.1016/0301-6226(91)90127-C:27:321-338.10.1016/0301-6226(91)90127-C.

33. Koch RM, L. A. Swiger, D. Chambers, Gregory. KE. 1963. Efficiency of Feed Use in Beef Cattle. Journal of Animal Science 10.2527/jas1963.222486x:22:486–494.doi.org/10.2527/jas1963.222486x.

34. Cantalapiedra-Hijar G, Abo-Ismail M, Carstens GE, Guan LL, Hegarty R, Kenny DA, McGee M, Plastow G, Relling A, I. O-M. 2018. Review: Biological determinants of between-animal variation in feed efficiency of growing beef cattle. . Animal doi:10.1017/S1751731118001489.:12(s2):s321-s335.10.1017/S1751731118001489.

35. Bergamaschi M, Tiezzi F, Howard J, Huang YJ, Gray KA, Schillebeeckx C, McNulty NP, C. M. 2020. Gut microbiome composition differences among breeds impact feed efficiency in swine. Microbiome. . doi:10.1186/s40168-020-00888-9. :8(1):110.10.1186/s40168-020-00888-9. .

36. Zhou Q, Lan F, Gu S, Li G, Wu G, Yan Y, Li X, Jin J, Wen C, Sun C, N. Y. 2023. Genetic and microbiome analysis of feed efficiency in laying hens. . Poult Sci doi:10.1016/j.psj.2022.102393.:102(4):102393.10.1016/j.psj.2022.102393.

37. McLoughlin S, Spillane C, Claffey N, Smith PE, O’Rourke T, Diskin MG, SM. W. 2020. Rumen Microbiome Composition Is Altered in Sheep Divergent in Feed Efficiency. Front Microbiol doi:10.3389/fmicb.2020.01981.:11:1981.10.3389/fmicb.2020.01981.

38. Li F, LL. G. 2017. Metatranscriptomic Profiling Reveals Linkages between the Active Rumen Microbiome and Feed Efficiency in Beef Cattle. Appl Environ Microbiol doi:10.1128/AEM.00061-17.:17;83(9):e00061-17.10.1128/AEM.00061-17.

39. Wensel CR, Pluznick JL, Salzberg SL, CL. S. 2022. Next-generation sequencing: insights to advance clinical investigations of the microbiome. J Clin Invest doi:10.1172/JCI154944.:132(7):e154944.10.1172/JCI154944.

40. Odom AR, Faits T, Castro-Nallar E, Crandall KA, WE. J. 2023. Metagenomic profiling pipelines improve taxonomic classification for 16S amplicon sequencing data. Sci Rep doi:10.1038/s41598-023-40799-x.:13(1):13957. .10.1038/s41598-023-40799-x.

41. Liu T, Wang Q, Gao C, Long S, He T, Wu Z, Chen Z. 2023. Drinking Warm Water Promotes Performance by Regulating Ruminal Microbial Composition and Serum Metabolites in Yak Calves. Microorganisms 11.10.3390/microorganisms11082092.

42. Trefts E, Gannon M, DH W. 2017. The liver. Curr Biol doi:10.1016/j.cub.2017.09.019:27(21):R1147-R1151.10.1016/j.cub.2017.09.019.

43. JY C. 2013 Bile acid metabolism and signaling. . Compr Physiol doi:10.1002/cphy.c120023:3(3):1191-212.10.1002/cphy.c120023.

44. Spector AA, Hoak JC, Fry GL, Denning GM, Stoll LL, JB S. 1980. Effect of fatty acid modification on prostacyclin production by cultured human endothelial cells. J Clin Invest doi:10.1172/JCI109752.:65(5):1003-12.10.1172/JCI109752.

45. S H. 1983. Leukotrienes. Annu Rev Biochem doi:10.1146/annurev.bi.52.070183.002035.:52:355-77.10.1146/annurev.bi.52.070183.002035.

46. Higgs EA, Moncada S, JR V. 1986. Prostaglandins and thromboxanes from fatty acids. . Prog Lipid Res doi:10.1016/0163-7827(86)90005-6.:25(1-4):5-11.10.1016/0163-7827(86)90005-6.

47. Tvrzicka E, Kremmyda LS, Stankova B, A. Z. 2011. Fatty acids as biocompounds: their role in human metabolism, health and disease--a review. Part 1: classification, dietary sources and biological functions. Biomed Pap Med Fac Univ Palacky Olomouc Czech Repub doi:10.5507/bp.2011.038.:155(2):117-30.10.5507/bp.2011.038.

48. Friedman MI, Harris RB, Ji H, Ramirez I, MG. T. 1999. Fatty acid oxidation affects food intake by altering hepatic energy status. Am J Physiol 4:R1046–53.10.1152/ajpregu.1999.276.4.R1046.

49. Rouzer CA, LJ M. 2009. Cyclooxygenases: structural and functional insights. . J Lipid Res Suppl:S29–34.10.1194/jlr.R800042-JLR200.

50. Cui J, J J. 2021. Natural COX-2 Inhibitors as Promising Anti-inflammatory Agents: An Update. Curr Med Chem 18:3622–3646.10.2174/0929867327999200917150939.

51. Francés DE, Motiño O, Agrá N, González-Rodríguez Á, Fernández-Álvarez A, Cucarella C, Mayoral R, Castro-Sánchez L, García-Casarrubios E, Boscá L, Carnovale CE, Casado M, Valverde ÁM, P. M-S. 2015. Hepatic cyclooxygenase-2 expression protects against diet-induced steatosis, obesity, and insulin resistance. Diabetes 5:1522–31.10.2337/db14-0979.

52. Hardy KD, Cox BE, Milne GL, Yin H, II RL. 2011. Nonenzymatic free radical-catalyzed generation of 15-deoxy-Δ(12,14)-prostaglandin J₂-like compounds (deoxy-J₂-isoprostanes) in vivo. J Lipid Res 1:113–24.10.1194/jlr.M010264. .

53. Korbecki J, Baranowska-Bosiacka I, Gutowska I, D. C. 2014. Cyclooxygenase pathways. . Cyclooxygenase pathways 61(4):639–49.

54. Metzler-Zebeli B, Siegerstetter S-C, Magowan E, Lawlor P, O’Connell N, Zebeli Q. 2019. Feed Restriction Reveals Distinct Serum Metabolome Profiles in Chickens Divergent in Feed Efficiency Traits. Metabolites 9.10.3390/metabo9020038.

